# The Correlation between MicroRNA-199a and White Adipose Tissue in C57/BL6J Mice with High-Fat Diet

**DOI:** 10.1101/167783

**Authors:** Dan Liu, Xia Wang, Xinying Lin, Baihui Zhang, Shue Wang, Wei Bao

## Abstract

Understanding is emerging about microRNAs as mediators in the regulation of white adipose tissue (WAT) and obesity. The expression level of miR-199a in mice was investigated to test our hypothesis: miR-199a might be related to fat accumulation and try to find its target mRNA, which perhaps could propose strategies with a therapeutic potential affecting the fat storage. C57/BL6J mice were randomly assigned to either a control group or an obesity model group (*n*=10 in both groups). Control mice were fed a normal diet (fat: 10 kcal %) *ad libitum* for 12 weeks, and model mice were fed a high-fat diet (fat: 30 kcal %) *ad libitum* for 12 weeks to induce obesity. At the end of the experiment, body fat mass and the free fatty acids (FFAs) and triglycerides (TGs) in WAT were tested. Fat cell size was measured by hematoxylin-eosin (H&E) staining method. The fat mass of the model group was higher than that of the control group (*P*<0.05). In addition, the concentrations of the FFAs and TGs were higher (*P*<0.05) and the adipocyte count was lower (*P*<0.05) in the model group. We tested the expression levels of miR-199a and key adipogenic transcription factors, including peroxisome proliferator activated receptor gamma2 (PPARγ2), CCAAT/enhancer binding proteins alpha (C/EBPα), adipocyte fatty acid-binding protein (aP2), and sterol regulatory element binding protein-1c (SREBP-1c). Up-regulated expression of miR-199a was observed in model group. Increased levels of miR-199a was accompanied by high expression levels of SREBP-1c. We found that the 3’-UTR of SREBP-1c mRNA has a predicted binding site for miR-199a. Based on the current discoveries, we suggest that miR-199a may exert its action by binding to its target mRNA and cooperate with SREBP-1c to induce obesity. Therefore, if the predicted binding site is confirmed by further research, miR-199a may have therapeutic potential for obesity.

**Abbreviations:** WAT, white adipose tissue; PPARγ2, peroxisome proliferator, activated receptor γ2; C/EBP αCCAAT/enhancer binding proteins α; aP2, adipocyte fatty acid-binding protein; SREBP-1c, sterol regulatory element binding protein-1c; HFD, high-fat diet.

## Introduction

Obesity is closely linked to metabolic syndrome, and it is a risk factor for various metabolic diseases, including type 2 diabetes, hypertension, hyperlipidemia and atherosclerosis(Kadakia, Fox *et al.* 2011; Kahn and Flier 2000; Malik, Willett *et al.* 2013). Excessive storage of white adipose tissue (WAT) is the main manifestation of obesity. WAT has been characterized as an endocrine organ that participates in energy metabolism (Alexander, Lodish *et al.* 2011). Although WAT is important in the regulation of metabolism, excessive fat accumulation in WAT is associated with metabolic syndrome(Wajchenberg 2000). Several key adipocytokines (e.g., adipocyte fatty acid-binding protein [aP2], CCAAT/enhancer binding proteins alpha [C/EBPα], peroxisome proliferator activated receptor gamma2 [PPARγ2], and sterol regulatory element binding protein-1c [SREBP-1c]) play a significant role in the regulation of fat storage (Chung, Kim *et al.* 2016; Yang, Vought *et al.* 2006; Zuo, Qiang *et al.* 2006), which in turn will affects the normal function of WAT.

MicroRNAs (miRNAs) are highly conserved, single-stranded non-coding RNAs(Ambros 2004). At present, numerous studies have confirmed that miRNAs are involved in the regulation of various human tumors, such as ovarian cancer, colorectal cancer, and others (Du and Sha 2017; Han, Zhao *et al.* 2017). Accumulating evidence has recently shown that miRNAs participate in the regulation of adipose tissue function, affecting adipose tissue metabolism and related diseases (Seton-Rogers 2012; van Rooij and Olson 2012). Many miRNAs are dysregulated in the WAT of obese animals and human subjects, potentially contributing to the pathogenesis of obesity-associated complications (Alexander, Lodish *et al.* 2011; Hilton, Neville *et al.* 2013; Peng, Yu *et al.* 2014). In addition, we also want to know the relationship between miRNAs and key adipocytokines, and whether their interaction leads to adipose tissue dysfunction. Despite wide study in the past decade, knowledge of the functional role between miRNAs and key adipocytokines remains limited. Further exploration of more miRNAs is required. Identification and exploration of more obesity-related miRNAs will be helpful for better understanding of fat accumulation and WAT dysfunction.

Among the many miRNAs that have been identified in humans, we focus on miR-199a because of its potential effect on obesity. Previous studies showed miR-199a was highly expressed in 3T3-L1 preadipocytes (Kajimoto, Naraba *et al.* 2006), and subcutaneous adipose tissue from piglets were shown to have a higher level of miR-199a(Shi, Li *et al.* 2014). Several studies have reported that miR-199a participates in the regulation of adipogenesis (Gu, You *et al.* 2016; Song, Gao *et al.* 2014), which indicates that miR-199a may play an important regulatory role in obesity. However, there is currently no report on the relationship between miR-199a and obesity.

In the present study, an obesity model was established in C57/BL6J mice. Free fatty acids (FFAs), triglycerides (TGs), adipocyte counts, and fat mass were observed among the obese and non-obese groups. The expression level of miR-199a in mice was investigated to test our hypothesis that miR-199a may be associated with fat accumulation. Furthermore, the study aimed to establish its target mRNA, which may propose strategies with a therapeutic potential that affect fat storage.

## Materials and methods

### Animals

All experimental protocols were approved by the Shandong University Institutional Animal Ethics Committee, and all procedures were performed in accordance with ethical standards. Male C57/BL6J mice (n=20) were obtained from PLA Nanjing Military Medical College (Nanjing, China). The weight of mice were 18~22g (4~6 weeks), and housing conditions was as follows: environment temperature 22~25℃, relative humidity 55%~65%, 12h light and 12 darkness cycle. The C57/BL6J male mice were randomly assigned to one of two groups (n=10 per group for this pilot study). In the control group, mice were fed a normal diet (10% kcal from fat) ad libitum for 12 weeks. In the model group, mice were fed a high-fat diet (30% kcal from fat) ad libitum for 12 weeks to induce obesity. Weight measurements were performed weekly. At the end of the experiment, adipose tissue mass was scanned in vivo using digital dual-energy X-ray (DEXA) scanners (Norland at Swissray. Fort Atkinson, WI, USA). After all of the mice were euthanized, WAT samples were dissected to detect associated indicators of obesity.

### Body fat mass

Fat mass was measured using digital DEXA scanners. DEXA data were processed for study of morphometry. Mice were anesthetized for ~1 h by injection of pentobarbital sodium (30 mg/kg bodyweight). The anesthetized mice were individually placed on a foam board and the scanning arm of the DEXA scanner assessed the fat mass (in grams) from neck to tail. After several minutes, the fat mass was recorded (Fig. 1).

**Figure 1.**
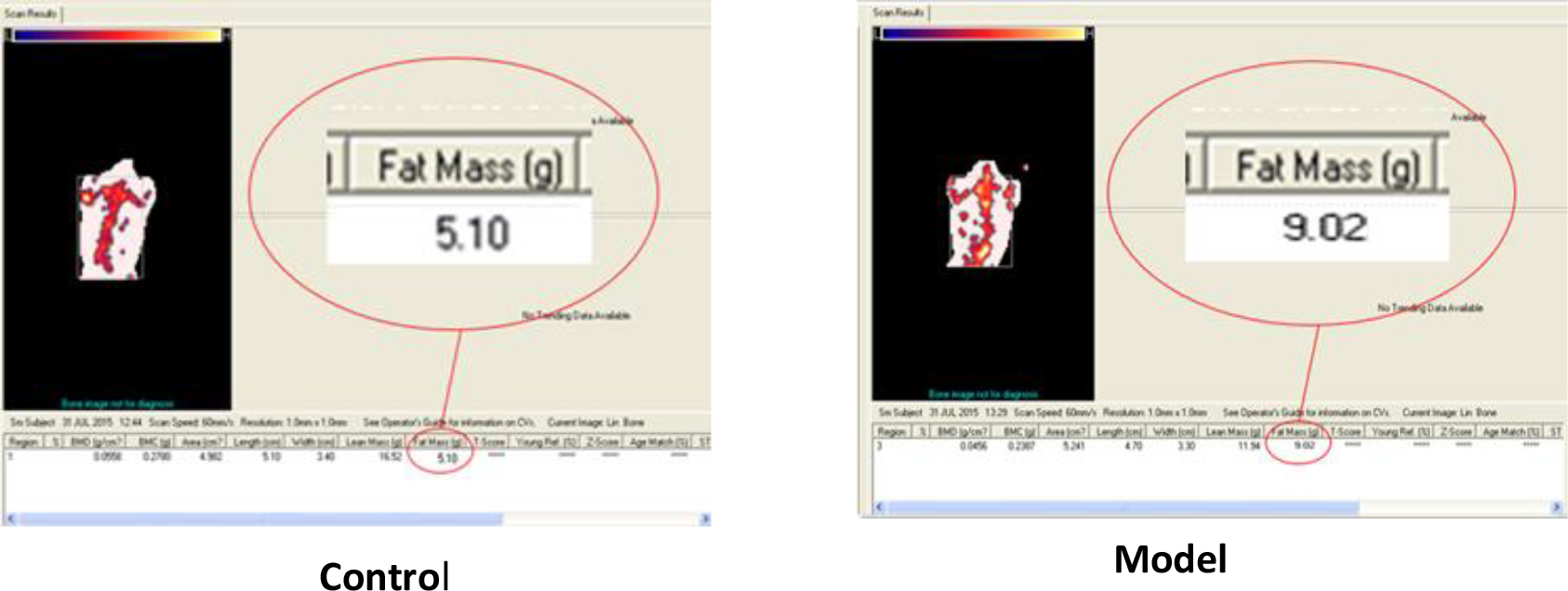
Body fat mass (from neck to tail) were scanned and tested by using digital DEXA scanners to process for morphometric studies. The fat mass of model group is higher than that of control group.

### FFAs and TGs levels in WAT

Subcutaneous adipose tissue and visceral adipose tissue were dissected, weighed, and stored at −80°C. Lipids from subcutaneous adipose tissue were extracted by the Folch method as previously described (Breil, Abert Vian *et al.* 2017), and the levels of FFAs and TGs were measured with the use of enzyme-linked immunoassays (Shanghai Tongwei reagent biological technology Co., Ltd., China) and the kit catalogue numbers were TWp002083 and TWp003393. The concentrations of FFAs and TGs were calculated based on a standard curve, which is made standard concentration (μmol/L) as abscissas and optical density (OD) value (lg (1/trans)) as ordinate.

### Adipocyte size measurement

Adipose tissue was stained with hematoxylin-eosin (H&E), and cells were counted and measured under the light microscope (×200 X, scale bar=50μm).

### Quantitative real-time polymerase chain reaction (qRT-PCR)

Total RNA was extracted from adipose tissue, which was isolated using an RNeasy Lipid Tissue Mini kit (Qiagen GmbH, Hilden, Germany) and stored at −80°C.The purity, concentration, and integrity of the RNA were analyzed by micro-ultraviolet spectrophotometry (Thermo Scientific NanoDrop 2000/2000c, USA) and agarose gel electrophoresis. Total RNA was reverse-transcribed with the ReverTra Ace qPCRRT kit (TOYOBO life science Co., Ltd., Japan) to obtain total cDNA. qRT-PCR was performed with SYBR^®^ Green qPCR Master Mix (Shanghai TOYOBO biological technology Co.,Ltd., China) in a LightCycler^®^ 480 system (Roche Group, Switzerland). The expression level of each mRNA was normalized to β-actin. SYBR Green fluorescence quantification method was used and thermocycling was 40 Ct. The primer sequences and cycling conditions used are listed in Table I.

### Statistical analyses

We used SPSS 22.0 software to analyze the data. Results are expressed as medians (P25, P75) or means ± SD according to normality test, which correspondingly used two independent-sample nonparametric tests or t-test to compare the medians or means between the two groups. Correlational analyses were calculated and are shown with correlation coefficient. A *P*-value <0.05 was considered statistically significant.

## Results

### Changes in lipid metabolism induced by high-fat diet

The body fat mass (from neck to tail) was determined using DEXA scanners (Fig. 1). The fat mass of the model group was higher than that of the control group (Table II; P=0.009). The H&E-stained adipocytes were measured under the light microscope (magnification, x200; Fig. 2 and Table II), and the number of adipocytes in the WAT samples was counted per unit area. Under a magnified visual field (x200) of the light microscope, an average number of 236 adipocytes were counted in the control group and 200 in the model group. The adipocyte count was lower in the model group than in the control group (Table II; P=0.012). The FFA and TG levels in the WAT are presented in Table II. The concentrations of FFA (Table II; P<0.05) and TG (Table II; P<0.05) were higher in the model group than in the control group.

**Figure 2.**
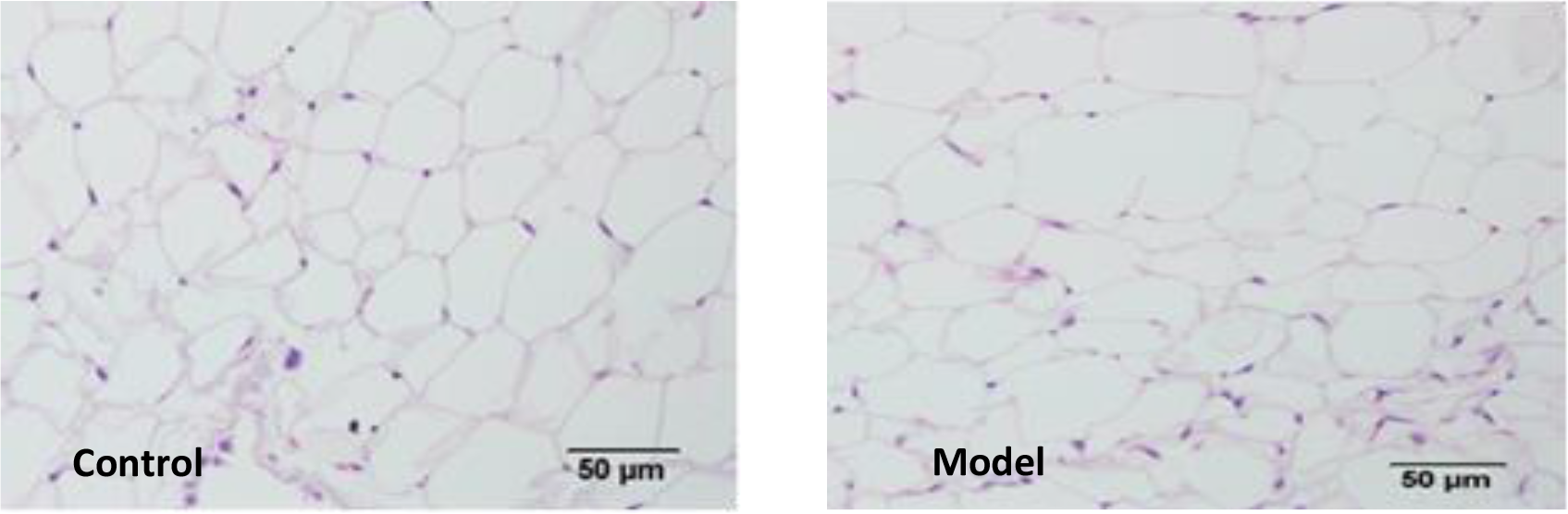
Adipocytes were stained with hematoxylin and eosin, and counted under the light microscope (magnification, x200; scale bar=50 μm). An average number of 236 and 200 adipocytes were counted in the control and model groups, respectively (Table II).

### Changes in the expression levels of miR-199a, aP2, C/EBPα, PPARγ2, and SREBP-1c

The miR-199a expression level in WAT was measured using RT-qPCR (Table III). The miR-199a expression level was upregulated in the model group and was higher when compared with the control group (Table III; P=0.037

To further confirm the role of miR-199a in obesity, the expression levels of four key adipogenic transcription factors, including PPARγ2, C/EBPα, aP2, and SREBP-1c were evaluated. The expression levels of aP2, C/EBPα, PPARγ2 and SREBP-1c were higher in the model group than in the control group. Aside from the expression level of C/EBPα (Table III; P=0.246), the increased expression levels of the three other indicators were statistically significant (Table III; P=0.021, 0.013 and 0.035).

Correlation between miR-199a and aP2, C/EBPα, PPARγ2, and SREBP-1c. The correlation (r) between miR-199a and the four key adipogenic transcription factors is presented in Table IV. SREBP-1c expression levels were correlated with the miR-199a expression level (P=0.002). No statistically significant correlation was observed between miR-199a and aP2, C/EBPα, or PPARγ2 expression levels (P >0.05). The functional link between these molecules was analyzed using the Miranda program. It was found that the 3’-UTR of SREBP-1c mRNA has a theoretical binding site for miR-199a. The miR-199a processing and recognition of the mRNA target sites are presented in Fig. 3.

**Figure 3.**
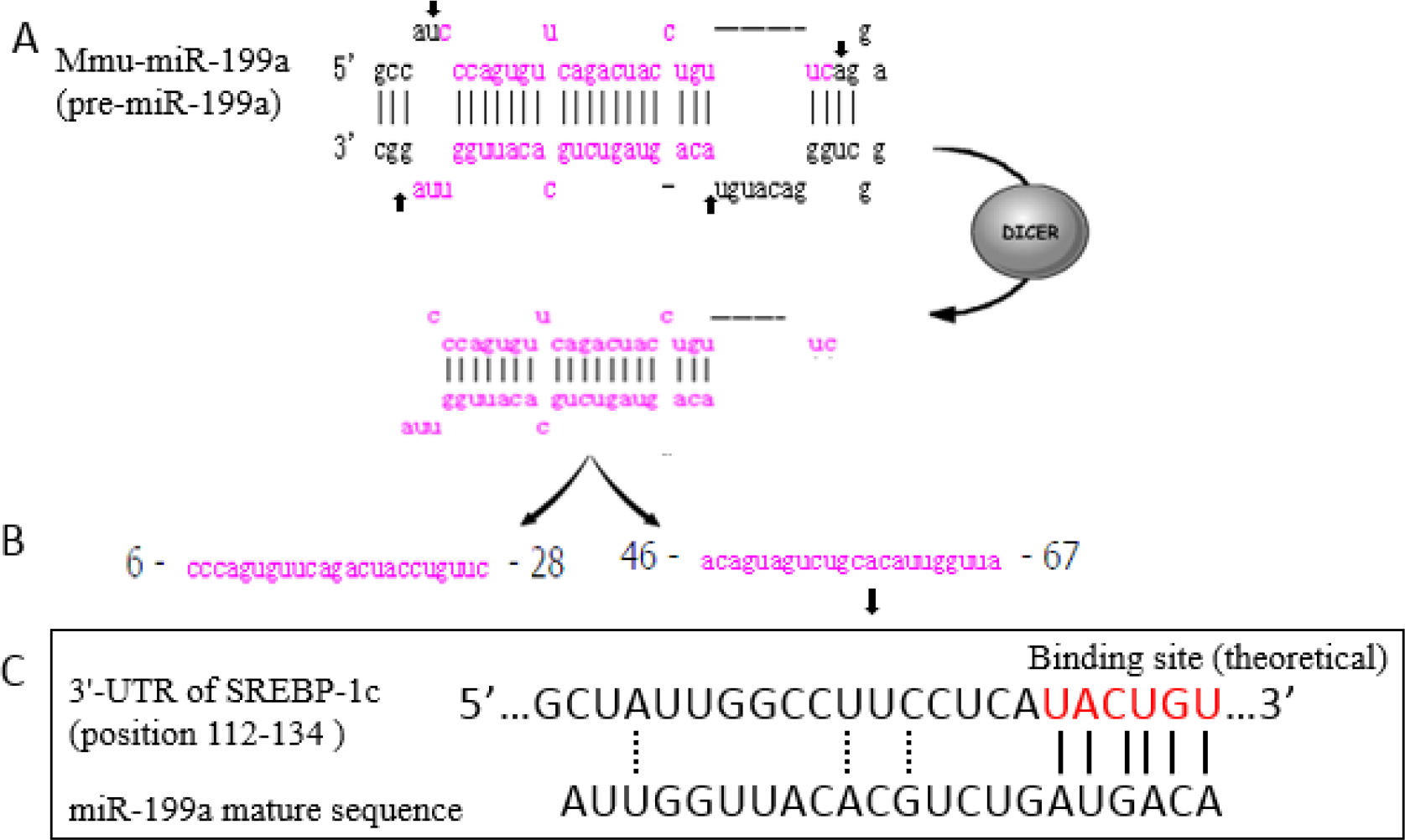
miR-199a processing and recognition of the mRNA target sites. (A) The hairpin structure of a pre-miRNA is processed into an miRNA duplex using the RNase Dicer via cleavage at various sites in the hairpin (indicated by the arrows). (B) The miRNA is unwound into two strands: miR-199a-3p (right) and miR-199a-5p (left). (C) The theoretical binding site (highlighted, vertical lines) is presented. The dotted lines mark interactions that influence and strengthen target recognition. miR, microRNA; UTR, untranslated region.

## Discussion

The results of our study showed increased expression level of miR-199a in the WAT of the model group. The expression levels of key transcriptional regulation factors related to adipocyte differentiation and fat accumulation, including aP2, PPARγ2, and SREBP-1c, were up-regulated in the WAT of the model group. Among them, SREBP-1c was correlated with the miR-199a expression level.

Obesity is an energy balance disorder that is characterized by increased lipid storage in adipocytes (hypertrophy) as well as an increased number of adipocytes (hyperplasia) (Wang, Tao *et al.* 2013). Adipocyte hypertrophy is the main reason of adult obesity. Generally speaking, it is not only caused by increased fatty acid synthesis from carbohydrates and fat intake by organs and low energy expenditure, but also by a malfunction of fat tissue. The typical diameter of an adipocytes is 0.3-0.9μm, but it can be about 20-fold larger in hypertrophic adipose tissue (Kim, Huh *et al.* 2015; Parlee, Lentz *et al.* 2014). In our study, a high-fat diet was used to create an obesity model. The adipocyte counts in the model group were lower than in the control group, but the fat mass was higher in the model group.

The function of WAT is to store excess energy in the form of TGs and to convert them into FFAs and glycerol to provide energy upon demand. When the excess calories overwhelm the storage capacity of adipocytes, excess fat is stored in the liver, leading to fat deposition and insulin resistance (Kershaw and Flier 2004). As an endocrine organ, WAT plays a crucial role in controlling whole body metabolism by secreting adipokines and storing FFAs(Choe, Huh *et al.* 2016), and increased WAT mass via hyperplasia and hypertrophy results in adipocyte dysfunction (Roberts, Hodson *et al.* 2009). In our study, the FFA and TG levels of the model group were higher than those of the control group. This finding implies a disorder in lipid metabolism in the hypertrophic adipose tissue.

MiRNAs, as transcription factors in the adipocyte differentiation process, were previously demonstrated to regulate adipogenesis and fat storage (Seton-Rogers 2012). A better understanding of the regulation of adipogenesis is crucial to the development of novel therapeutic strategies for obesity and its associated metabolic syndromes. Existing data demonstrated that miR-33 reduces the oxidation of fatty acids and inhibits the production of high-density lipoprotein (HDL), and it appears to be up-regulated in the individuals with obesity (Su, Zhang *et al.* 2017). In addition, the miR-222 expression level was found to be up-regulated in the serum of obese subjects, whereas miR-221 was down-regulated (Ortega, Mercader *et al.* 2013). The expression levels of both types of miRNAs are related to body mass index (BMI), waist circumference measurement, fat distribution, and HDL concentration (Ortega, Mercader *et al.* 2013). Many studies have also shown that miRNAs can be used as targets in the treatment of obesity and obesity-related chronic diseases (Alexander, Lodish *et al.* 2011; Kolfschoten, Roggli *et al.* 2009; Peng, Yu *et al.* 2014).

As shown in Figure 3, pre-miR-199a has a hairpin structure. This hairpin structure is processed into a miR-199a duplex by the RNase Dicer through cleavage at several sites in the hairpin. The miRNA is then unwound into two strands: miR-199a-3p (right) and miR-199a-5p (left). A previous study showed that miR-199a was highly expressed in 3T3-L1 preadipocytes (Kajimoto, Naraba *et al.* 2006), and another study demonstrated that the subcutaneous adipose tissue from piglets had a higher level of miR-199a(Shi, Li *et al.* 2014). Other several researches have shown that miR-199a participates in the regulation of adipogenesis (Gu, You *et al.* 2016; Song, Gao *et al.* 2014), suggesting that miR-199a may play an important regulatory role in obesity. However, the potential role of miR-199a in adipogenesis and hypertrophy of WAT has not yet been demonstrated in animal models or in human subjects. In our study, the result showed an increased expression level of miR-199a in the WAT of the model group. Based on these results, we hypothesized that the increased miR-199a expression was associated with lipid accumulation. We then conducted the experiments to test this hypothesis.

We also measured the expression levels of key transcriptional regulation factors that are related to adipocyte differentiation and fat storage. PPARγ, which is specifically expressed in WAT, is involved in lipid formation in mature adipocytes (White and Stephens 2010). C/EBPα also stimulates adipogenesis and works together with PPARγ in the process of adipocyte differentiation (Zuo, Qiang *et al.* 2006). In addition, other transcription factors, including aP2 and SREBP-1c are related with to fatty acid metabolism and glucose metabolism (Kolehmainen, Vidal *et al.* 2001; Kralisch and Fasshauer 2013; Wang, Kouri *et al.* 2005). Our results showed that the expression levels of aP2, PPARγ2, and SREBP-1c were up-regulated in the WAT of the model group. The results were consistent with previous reports (White and Stephens 2010). Moreover, SREBP-1c expression was consistent with the miR-199a expression level (Table IV), implying a potential mechanism between obesity and the miR-199a. In the control group, SREBP-1c expression level was low. When induced by a high-fat diet, the expression increased significantly in line with miR-199a expression. It showed that miR-199a may be relevant to obesity induced by a high-fat diet and the potential mechanism might be associated with SREBP-1c. We analyzed the functional link of these molecules by Miranda programs (http://www.microrna.org/microrna/home.do) and found that the 3’-UTR of SREBP-1c mRNA has a theoretical binding site for miR-199a. And the miR-199a processing and recognition of the mRNA target sites was showed in figure 3C. Based on our findings, we speculate that miR-199a may exert its action by binding to its target mRNA and cooperate with SREBP-1c to induce obesity. As reported in the literature, miRNAs function is by partially pairing to sequences located in the 3’UTR of target mRNA (Abelson, Kwan *et al.* 2005). If the predicted binding site for miR-199a in the 3’UTR of SREBP-1c is confirmed in further research, therapeutic strategies affecting the function of miR-199a may be possible (e.g., chemically modified complementary inhibitors). However, future research would first need to verify the direct role of miR-199a by using gene silencing or knockout technology specifically in WAT.

In conclusion, the present study showed that miR-199a expression is increased by a high-fat diet, and a higher level of miR-199a is correlated with increased expression of SREBP-1c. We analyzed the functional link of these molecules and found that the 3’-UTR of SREBP-1c mRNA has a theoretical binding site for miR-199a. Further research is needed to confirm our speculation. If the predicted binding site is confirmed in future research, potential therapeutic strategies affecting the function of miR-199a may be proposed for obesity and related-diseases.

## List of abbreviations

White adipose tissue (WAT)

Peroxisome proliferator activated receptor gamma2 (PPARγ2)

CCAAT/enhancer binding proteins alpha (C/EBPα)

Adipocyte fatty acid-binding protein (aP2)

Sterol regulatory element binding protein-1c (SREBP-1c)

High-fat diet (HFD)

## Acknowledgments

The authors declare no conflicts of interest that would prejudice the impartiality of this scientific work. This work was sponsored by National Natural Science Foundation (NSFC 81370966) of China.

Table I List of gene specific primers for real-time PCR

Table II Male C57/BL6J mice fat metabolism parameters (control, normal diet; model, high-fat diet), Medians (P25, P75) or Mean ((mean-SD)-(mean + SD))

Table III The relative expression level (2^-ΔΔCT^) of miR-199a, aP2, C/EBPα, PPARγ2 and SREBP-1c in two groups, Medians (P25, P75) or Mean ((mean-SD)-(mean +SD)) Table IV The correlation(*r*) between miR-199a and aP2, C/EBPα, PPARγ2 and SREBP-1c

